# Accelerated Simulation of Large Reaction Systems Using a Constraint-Based Algorithm

**DOI:** 10.1101/2020.10.31.362442

**Authors:** Paulo E. P. Burke, Luciano da F. Costa

## Abstract

Simulation of reaction systems has been employed along decades for a better understanding of such systems. However, the ever-growing gathering of biological data implied in larger and more complex models that are computationally challenging for current discrete-stochastic simulation methods. In this work, we propose a constraint-based algorithm to simulate such reaction systems, called the Constraint-Based Simulation Algorithm (CBSA). The main advantage of the proposed method is that it is intrinsically parallelizable, thus being able to be implemented in GPGPU architectures. We show through examples that our method can provide valid solutions when compared to the well-known Stochastic Simulation Algorithm (SSA). An analysis of computational efficiency showed that the CBSA tend to outperform other considered methods when dealing with a high number of molecules and reaction channels. Therefore, we believe that the proposed method constitutes an interesting alternative when simulating large chemical reaction systems.

## 1 Introduction

Computational models of biochemical reactions have proven to be useful tools for a better understanding of molecular systems [1, 2], as well as in optimizing experimental research [3, 4]. As a consequence of ever increasing data availability, these models are significantly growing in size and complexity toward the scale of entire cells [5, 6, 7].

Simulations of such models can be carried out by applying among a choice of many available methods (e.g., Ordinary Differential Equations, Stochastic Simulation Algorithms, and Dynamic Flux Balance Analysis) [8, 9]. However, each simulation method has its own advantages and limitations, which influence their choice concerning specific applications.

Complex reaction systems, such as biochemical interactions in a cellular environment, have some characteristics that are important to consider when modeling and simulating it. One of these characteristics is that complex biochemical systems often involve reactions taking place at significantly different time-scales. Also, the number of copies of molecular species can vary from a single to thousands of molecules [10, 11].

Another important characteristic is that the number of copies of a given molecular species at any time takes a discrete value. However, when the system involves a sufficiently high number of copies for all species involved, it is reasonable to treat this quantity as a continuous variable [12]. With this assumption, the most straightforward approach to simulate such systems is to solve a set of coupled, first-order, ordinary differential equations called Reaction-Rate Equations (RE) [13, 14, 15]. Other approaches have been developed for the case when the rate-constants are not available. It is the case of the Dynamic Flux Balance Analysis [8, 16]. Despite the computational efficiency of these methods for the case of many molecules, when it comes to smaller numbers, the results can be unsatisfactory in the sense that they will provide real-valued approximations of quantities that are in fact discrete-valued. Also, these methods provide deterministic solutions, while reactions are better described as stochastic processes [17, 18].

A physical consequence of a small number of copies of a given molecule in a biochemical system is that the reactions involving it will have their rate highly influenced by thermal fluctuations and molecular crowding. A better description of these reaction rate fluctuations can be achieved through stochastic formulations of the system. Then, the evolution of molecular counts would be a consequence of the reaction occurrences following a given probability rather than a rate. The single-valued function that gives the probability of finding a particular set of molecular counts at a given time is called the Reaction Master Equation (RME) [19]. Its behavior in time can be formulated as a Markovian random walk in RME space. Although it is often impossible to solve an RME analytically, the most frequently used method being equivalent to numerically solving it is the Stochastic Simulation Algorithm (SSA), also known as the Gillespie Algorithm [20, 21]. Even though SSA can adequately account for the inherent discreteness and stochasticity of molecular interactions, it typically implies a high computational cost. Some faster methods were derived from approximations of SSA, such as the *τ* Leaping method [22, 23]. However, in such cases, the efficiency can still be limited when considering bigger, more realistic biochemical models.

So far, we pointed out some important characteristics shared by many complex biochemical systems, such as different scales of reaction rates and molecular concentrations, discrete-valued number of molecules, and random fluctuations in reaction rates. Therefore, an ideal simulation method for such systems should be able to encompass all these features while being computationally efficient. Here we present an alternative method, henceforth called Constraint-Based Simulation Algorithm (CBSA), to simulating reaction systems focused on computational scalability while considering the system discreteness, stochasticity, and possible differences in time-scales. We use a constraint-based formulation of the models following the well-established methodology applied to Flux Balance Analysis [24]. Then the simulation’s computation is entirely performed using vector and sparse-matrix operations, which are highly parallelizable, thus being able to take advantage of GP-GPU architectures.

Although we do not perform a rigorous proof of the method, we show by using theoretical models that the numerical solutions of CBSA can be very close to the ones provided by the Stochastic Simulation Algorithm. Using a model of diffusion in a discrete toroidal-lattice space, we compare the computational efficiency between CBSA and some SSA implementations. The CBSA tends to outperform other algorithms in computational time for the case of large systems as a consequence of its ability to be computed on GPUs.

This manuscript is organized as follows. First, we introduce the Stochastic Simulation Algorithm, the *τ*-Leaping method, and the mathematical formulation of constraint-based (CB) models. Next, we discuss the flux estimation on CB models and its extension in order to accommodate a broader range of reactions. In Section 5, we describe the so-proposed Constraint-Based Simulations Algorithm, followed by an example in which we try to demonstrate its correctness in Section 6. Given that computational scalability is one of our main goals, in Section 8, we include a comparison between different implementations of SSA methods and the CBSA regarding their computational efficiency on large chemical systems using a diffusion model. In the last section, we provide some concluding remarks on this work.

## 2 The Stochastic Simulation Algorithm

Since its publication by Daniel Gillespie in [20], the Stochastic Simulation Algorithm (SSA) has been the standard approach to simulate stochastic systems. In the most used form of this algorithm, called the *Direct Method*, a well-mixed solution where *M* molecular species can react through *R* reaction channels is assumed. Given an initial state *X*_0_ for the number of copies of each molecular species, we define a time-step *τ* where only one of the *R* possible reactions can occur. For each reaction channel *μ* in *R*, we must calculate its propensity such as:

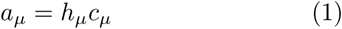

where *h*_*μ*_ is the number of possible combinations between the reactant molecules of *μ* in the current state of the system *X*(*t*). The *c*_*μ*_ is a reaction constant that can be understood as the probability of that reaction occurs in a period. It is related to the rate constant of the reaction *k*_*μ*_ and the volume of the system *V* as follows:

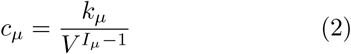

given that *I*_*μ*_ is the number of molecules reacting in *r*_*μ*_. The simulation step is then performed by selecting a pair of *μ* and *τ* such as:

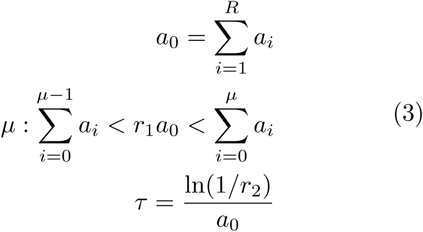

where *r*_1_ and *r*_2_ are random numbers drawn from a unitary uniform distribution. We then update the system state *X* by performing the reaction *μ* and then increase time *t* by *τ*. This procedure is repeated until a stopping criterion is reached.

Although SSA has been extensively tested and applied in the simulation of several stochastic systems, it tends to be a computationally expensive algorithm which may become prohibitive for very large systems [18]. There are some optimized formulations and approximation of the SSA aimed at improving computational performance [22]. In the following subsection, we present one of the most used approximation methods, the *τ*-Leaping method.

### 2.1 The *τ*-Leaping Method

The *τ*-Leaping method, also proposed by D. Gillespie [22], is an approximation of SSA which is considerably faster to compute. In this method, at each iteration, we leap in time by *τ* so that the changes in the propensity function *a* will be slight. Then, the number of occurrences *k*_*μ*_ in the time step *τ* for each reaction channel *μ* ∈ *R* will be sampled from the Poisson random distribution:

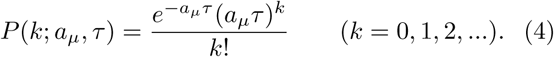

## 3 The Constraint-Based Modeling

Let *m* = {*m*_1_, *m*_2_, …, *m*_*M*_} be a set of *M* molecular species and *r* = {*r*_1_, *r*_2_, …, *r*_*R*_} be a set of *R* reactions where molecules in *m* play roles as reactants and/or products. The stoichiometry of each reaction, *i*.*e*. how much of each molecular species is consumed or produced, can be represented by means of a stoichiometric matrix *S*_*M×R*_. Every entry *S*_*ij*_ yields how much of the molecular species *i* is consumed (negatively signed) or produced (positively signed) by the reaction *j*. If we consider a vector *x* of length *M*, where *x*_*i*_ is the number of *m*_*i*_ molecules, homogeneously distributed in a given volume V, the variation of *x* along time can be written as:

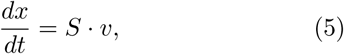

where *v* is a vector of fluxes and has length *R*. The flux *v*_*i*_ can be understood as how many times the reaction *i* occurs in a given period. The latter will henceforth be considered as 1 second.

Although *v* is, for now, an unknown variable, its values are constrained by well-known physical properties such as mass balance. Therefore, we can set the following constraints to the solution space:

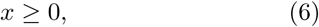

which assure that there are no negative number of molecules, and

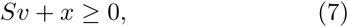

asserting that a given flux is valid if, and only if, it does not result in any negative number of molecules. An example of a solution space for two parallel reactions is depicted in Figure 1. The allowable space is upper-bounded the line that determines the mass-balance constraint.

**Figure 1:**
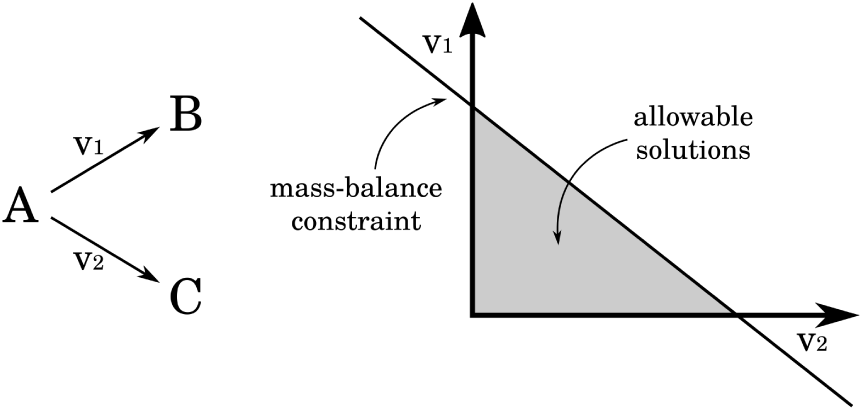
Example of a solution space for a system with two reactions. The flux for each reaction is represented in the axes. Both reactions share the same reactant A, thus, the combination of fluxes *v*_1_ and *v*_2_ must satisfy the mass-balance constraint.

## 4 Flux Estimation

The flux *v*_*i*_(*t*) of a reaction *r*_*i*_ in a given instant *t* can be understood as how many times the reaction *r*_*i*_ can occur along the next second. Therefore, the flux of a given reaction can be any function *g*(*x*) multiplied by the total number of combinations between the reactants. For the sake of simplicity, we will consider *g*(*x*) as a constant *c*_*i*_ for each reaction. Thus, following the same reasoning used in SSA to estimate *a*, we can write *v*_*i*_(*t* + Δ*t*) as follows:

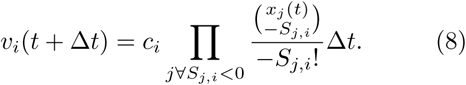

As we use the stoichiometry matrix to estimate *v*, the flux will be correct only if all reactants are different from products for all reactions. This is because *S* cannot yield information about the molecular species which counts does not change by the end of the reaction, even being necessary in the process. This is the case of catalysts, modifiers, or auto-catalytic reactions (*e*.*g*. A → A+A). Thus, to represent such cases, we will define a regulation matrix 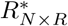 where 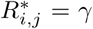 if *γm*_*i*_ molecules are needed in reaction *r*_*j*_. Then, we can rewrite Eq. 8 as:

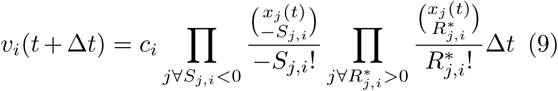

## 5 The Constraint-Based Simulation Algorithm – CBSA

In the last sections, we presented the most used algorithm applied to the stochastic simulation of chemical reactions called SSA, a mathematical formalism to describe such reactions and their behavior as a system of constraints, named Constraint-Based Modeling, and an expression to calculate fluxes in each reaction channel. Now, we will use these concepts to formulate a scalable algorithm that aims at simulating chemical reaction systems while considering their inherent discreteness and stochasticity.

Given an initial condition, each iteration of the algorithm is divided into five steps: 1) calculate the flux of each reaction based on the current system state; 2) add noise; 3) check if the calculated fluxes satisfy the mass-balance constraints (Eq. 7); 4) if mass-balance constraint not satisfied, find a valid set of fluxes; and 5) update the molecular counts. Figure 2 illustrates the algorithm scheme. These steps must be repeated until a given stopping criterion is reached, such as a maximum time.

**Figure 2:**
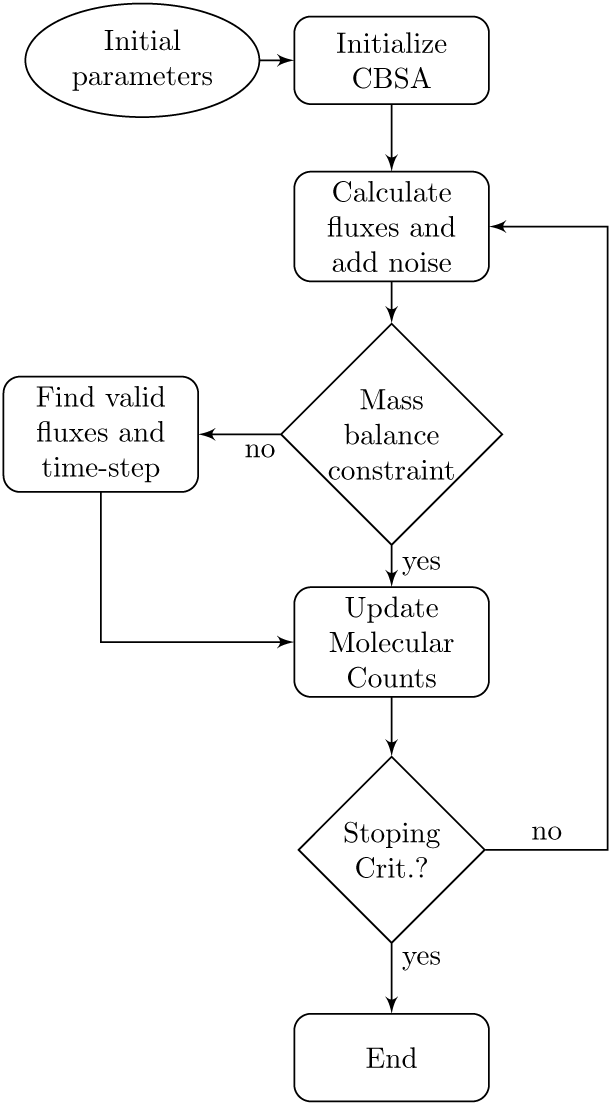
Flowchart of the Constraint-Based Simulation Algorithm.

The following paragraphs describe in detail each step of the algorithm.

### Initial Parameters

The chemical system is defined by the stoichiometric matrix *S*, which describes the reactants and products of every reaction, and the regulation matrix *R*^*\**^ that indicates other molecules necessary to given reactions but without being consumed nor produced in the process. Each reaction must also have an associated constant *c*, which is the fraction of all combinations between its reactant molecules that would indeed result in a reaction by the time of one second. Also, the initial state of the system *x*_0_ must be provided, defining the initial molecular count for every molecular species. Finally, we need to set a maximum time-step length Δ*t*_*max*_.

### Adding Noise to Fluxes

In order to account for fluctuations which might influence the velocity of reactions, specially in the case of lower molecular counts, we will add noise to the calculated flux on each step using the chemical Langevin Equation [25, 22]. This equation introduces a Gaussian white noise *η* in the flux *v* (Eq. 13) as:

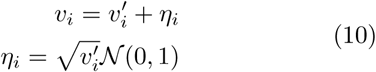

where 𝒩 (0, 1) is a random variable drawn from the normal distribution with mean zero, and standard deviation one. The variable 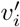 is equal to 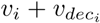 which will be introduced in the next paragraph.

### Discrete Flux Calculation

In order to account for the inherent discreteness of the system, the flux *v* must be always integer. However, Eqs. 9 and 10 can output decimal results. In order to keep *v* ∈ ℕ, we will separate the integer from the decimal part and store the later in *v*_*dec*_ for the next iteration. Then, *v*_*dec*_ and *v* at each iteration can be calculated using the following operations:

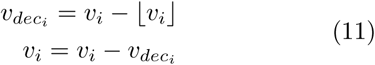

By storing the decimal part in 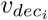, we assure that reactions in which *v*_*i*_ < 1 will occur at a delayed time when *v*_*i*_ ≥ 1.

### Finding a Valid Flux

Although each *v*_*i*_ is mostly valid for every single reaction, this might not hold when considering the whole system. To illustrate such a case, consider a system in which a given molecular species *m*_*k*_ is a reactant for many different reactions. Once the estimation of each *v*_*i*_ is independent of the others, the total amount of *m*_*k*_ required for all reactions might exceed the pool *x*_*k*_ and violate the mass-balance constraint from Eq. 7. It may also happen in the case of *c*_*i*_ >> 1. To avoid such condition, if *v* fails the mass-balance constraint, we may look for the maximal Δ*t* that assure non-negative values on *x*(*t* + Δ*t*). We do that by multiplying the current Δ*t* by a constant 0 < *α* < 1, recalculating the fluxes, updating the decimal part *v*_*dec*_, and testing the mass-balance constraint until a valid solution is found, as shown in Eq. 12.

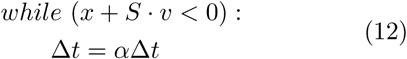

In the worst-case scenario, all *v*_*i*_ will reach zero, and another calculation step begins restoring Δ*t* to its initial value.

### Molecular Count Update

Having a discrete flux *v* passing the mass-balance constraint stated in Eq. 7, we simply update the molecular counts by solving Eq. 5 using Euler’s method. The update equation becomes:

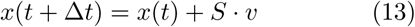

## 6 Biochemical System Simulation

In order to test the correctness of the solutions provided by the CBSA, we use here a reaction system that can demonstrate several of the important characteristics of real biochemical systems, such as different magnitudes of molecular counts and reaction velocities, and regulated reactions while demonstrating the method’s capabilities and limitations. Other examples can be found in the Supplementary Information.

Let us consider the reaction system {1} where a Transcription Factor (*Tf*) triggers the production of a protein *P* when active. The Transcription Factor *Tf* is activated and deactivated spontaneously by a slow reaction. The activated *Tf* triggers a swift production of hundreds of protein *P* copies, which are spontaneously degraded. With the system starting with only one *Tf* molecule, it should transit between its activated and inactivated states, triggering bursts on the production of *P* (Fig. 3a).

**Figure 3:**
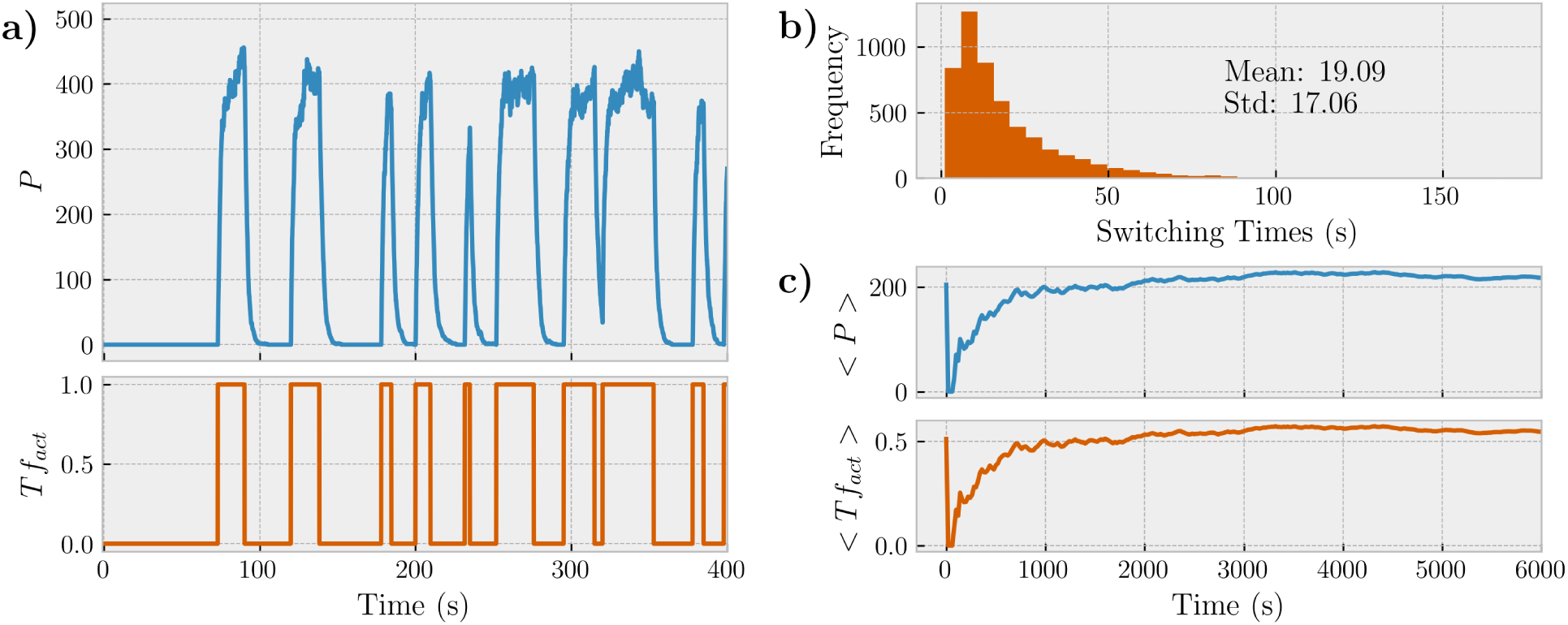
**(a)**Simulation of the chemical system described in {1} using the CBSA with *c*_1_, *c*_2_, *c*_3_, and *c*_4_ equals to 0.05, 0.05, 200, and 0.5 respectively. The initial state of *x* contains only one *Tf*_*inactive*_ molecule. Every time that *Tf*_*inactive*_ turns into *Tf*_*active*_, a burst in *P* is observed. **(b)** With *c*_1_ = *c*_2_ = 0.05, the *Tf* should transit between states every 20 seconds in average. We can see the distribution of switching times with mean 19.09 and standard deviation 17.06 obtained from a simulation of 100,000 seconds. **(c)** The cumulative mean of *Tf*_*active*_, named < *Tf*_*act*_ >, at a given time *t* corresponds to the mean value of *Tf*_*active*_ in the interval [0, *t*]. Once *c*_1_ = *c*_2_, they have the same transition probability, thus, < *Tf*_*act*_ > should converge to 0.5 with *t* → ∞. Likewise, the cumulative mean of *P* should converge to 200 once the number of *P* molecules peaks at around 400 when *Tf* is active, and it is approximately for half of the time. The parameter *α* was fixed as 0.5 for all CBSA simulations.

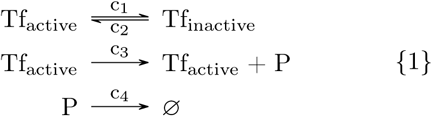

The simulation depicted in Figure 3a was performed using the CBSA. The biochemical system is represented employing the stoichiometric matrix *S* and the regulation matrix *R*^*\**^. As *S* can only represent the reaction’s final stoichiometry without accounting for intermediate molecules, we must represent the catalytic *Tf*_*active*_ molecule in the regulation matrix *R*^*\**^ as follows:

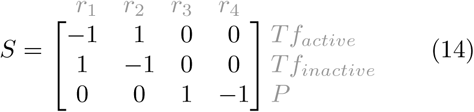

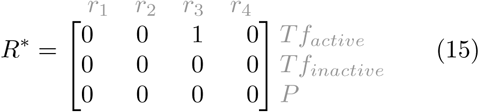

where *r*_1_ to *r*_4_ are the respective reaction channels with rate constants *c*_1_ to *c*_4_.

The times between activation and deactivation of *Tf* depend solely in the rate constants *c*_1_ and *c*_2_. Thus, we could estimate these times as 1*/c*_1_ and 1*/c*_2_. In the example depicted in Figure 3a we adopted *c*_1_ = *c*_2_ = 0.05 making the transition times between activate and inactivate states equals to 20 seconds. However, the system is stochastic and the observed transition times will compose a distribution rather than an exact value. In Figure 3b we can observe the distribution of the switching times between states of *Tf* with the mean close to the expected value of 20 seconds. Given any equal values for *c*_1_ and *c*_2_ and the initial condition with only one *Tf* molecule, we can estimate that the mean value for both activated and inactivated states should converge to 0.5 with *t* → ∞. This behavior is indeed observed in Figure 3c.

### 6.1 Validation Respectively to the Stochastic Simulation Algorithm

The SSA is already a well-established method to perform stochastic simulations of chemical systems. Thus, we use it as a gold-standard method in order to evaluate the correctness of the simulations performed using the proposed Constraint-Based Simulation Algorithm.

Regarding the reaction system described in {1}, we already observed agreement between its simulations using CBSA and theoretical mean values expected, as shown in Figure 3. Now, we shall compare these results with trajectories obtained from simulations using the original Stochastic Simulation Algorithm and its approximated form, the *τ*-leaping. Tough all methods are stochastic, it is not necessary to perform several runs of the same experiment to obtain statistics about the model because this is a particular oscillatory system and does not diverge at any point in time. Thus, we can make consistent statistics from the time-series with sufficiently long simulation time.

The first comparison we make regards the distribution of switching times between both states of *Tf*. In Figure 4a we can see that the distributions for the three considered methods have the same exponential-like shape with similar means and standard deviations. With the SSA and *τ*-leaping method, we obtained the mean switching times closer to the theoretical value. It is also observed that the CBSA allowed a lower standard deviation than the other methods. We believe this is a consequence of the constraints imposed on the model after the addition of noise, which can reject nonviable solutions. In this sense, the constraints may introduce a bias in the distribution from which the random numbers are drawn mainly affecting the standard deviation of reaction times.

**Figure 4:**
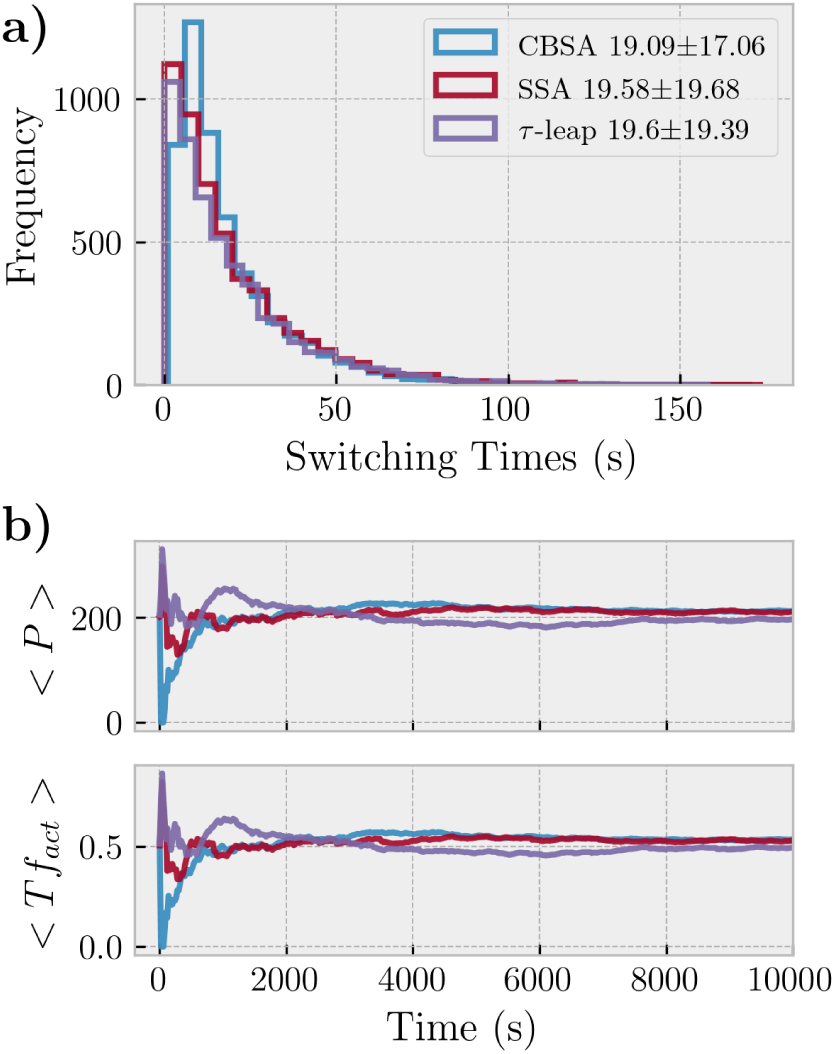
**(a)** Distribution of *Tf* switching times for the CBSA, SSA, and *τ*-leaping methods with a simulation time of 100,000 seconds. Means and standard deviations indicated in the legend. **(b)** The cumulative mean of *P* and *Tf*_*active*_ for the three methods.

A consequence of the lower standard deviation of reaction times can be observed in the time that the system takes to converge the mean number of molecules to the theoretical value. The mean values of both *P* and *Tf*_*active*_ converge faster to the expected values as depicted in Figure 4b. Another limitation of the methods is that it does not necessarily lead to a linear relationship between the time step adopted in the Euler integration and the error obtained when compared to the theoretical value. The main reason for this possible non-linear relationship between time step and error is that the noise *η*_*i*_ added to each flux *v*_*i*_ goes with the square-root of the flux. So, when *v*_*i*_ < 1, the smaller the value that *v*_*i*_ takes, the greater is the noise *η*_*i*_ introduced. Thus, by reducing the integration step Δ*t* we increase the chance o obtaining a flux *v*_*i*_ < 1 and the inverted noise proportion.

Despite the differences in the standard deviation of reaction velocities, all methods agreed regarding the qualitative behavior of the system and molecular counts. Other examples can be found in the Supplementary Information where similar conclusions are obtained. We also discuss other aspects of the algorithm such as the influence of Δ*t* and *α* in the results, relative errors, and in the number of steps to achieve the simulation time.

## 7 Accelerated Computing of The CBSA

One of the main constraints of the Stochastic Simulation Algorithm is its computational cost. The algorithm performs only one reaction for each time step and, depending on the size of the system and the rate constants, it may demand a very large number of steps to achieve the desired simulation time. The *τ*-leaping approximation and some variations of it can reduce the number of steps in some cases, usually being adopted for bigger chemical systems.

Another limitation of SSA is that the way the algorithm is designed makes it difficult to be implemented for parallel environments. Although there are implementations of the SSA to be computed using many computer cores, or even using General-Purpose Graphics Processing Units (GP-GPUs) [26, 27, 28], they can only accelerate replicates of the same model, reducing the time to compute several trajectories, but still limited regarding the size of the chemical system. There are also implementations for computing the SSA in a discretized space [29] where the subsystems in each subspace can be performed in parallel.

The CBSA is almost entirely based on vector and matrix operations, being fast to compute and easy to implement. Also, considering that the matrices *S* and *R*^*\**^ are often highly sparse, a more sophisticated implementation can take advantage of this feature to further enhance computations regarding both processing time and memory usage. A description of the sparse implementation can be found in the Supplementary Information.

Parallel computing was a design principle when developing the reported method. All operations in the CBSA can be computed in parallel thanks to the relative independence of the operations through the vectors and matrices. Therefore, we could take advantage of the massive computing power of GP-GPUs to accelerate individual simulations of large chemical systems.

## 8 Computing Time Benchmark

One of the goals of the present work is to develop a simulation method that is computationally scalable to a variety of chemical system sizes. By size, we mean both the total number of molecules and the number of reactions. We shall then compare, for some cases, the CBSA’s computational expense against the SSA and some of its implementations, and also an ODE solver. ^1^ For the SSA, we use the implementations from three Python libraries: StochPy^2^ [30], STEPS^3^ [31], and GillesPy2^4^ [32]. The ODE implementation is part of the latter library. We use pyOpenCL^5^ to distribute computations of the CBSA on GPUs.

### 8.1 Benchmark Model

We use as a benchmark model a diffusion process through a discrete toroidal square-lattice space of length *L*, as depicted in Fig. 5a. In this model, reactions will represent transitions between neighbor sub-spaces with a diffusion coefficient *k*_*diff*_. There-fore, a system with a space of length *L* yield 4*L*^2^ reactions and *L*^2^ molecular species. The system will start with an initial amount of molecules in only one of the sub-spaces and they will diffuse with a rate *k*_*d*_.

**Figure 5:**
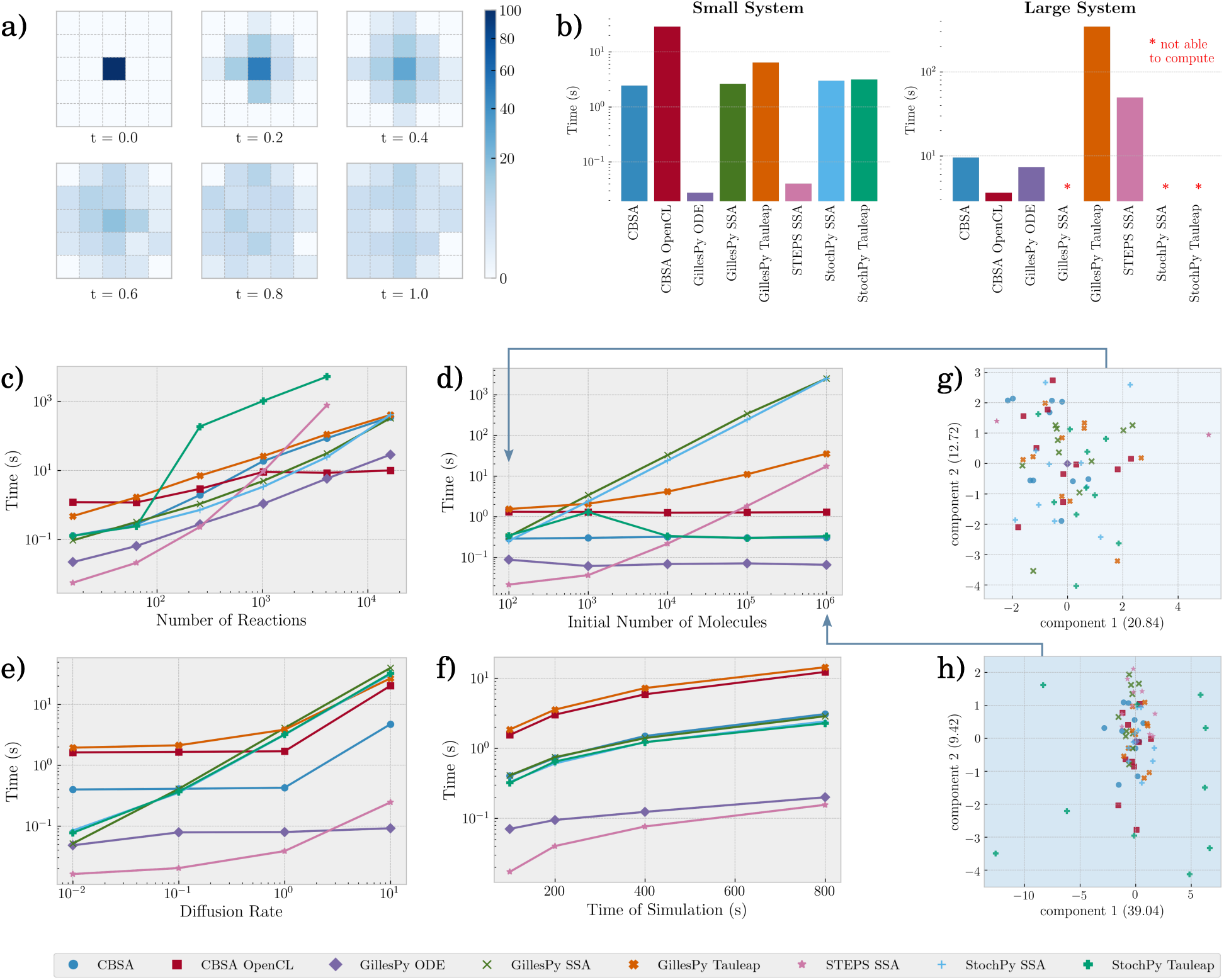
Benchmark using a diffusion model though a square-lattice discretized space with periodic boundary conditions. **(a)** Simulation of the system with length *L* = 5 and 100 initial molecules using CBSA. Each plot depicts the distribution of the diffused molecules in the grid for the times *t* = {0.0, 0.2, 0.4, 0.6, 0.8, 1.0}. **(b)** Simulation times in logarithmic scale for a small and a large system. The parallel implementation of the CBSA is denoted as “CBSA OpenCL”. Considering that the CBSA is designed to simulate large systems, we selected two different scenarios where we estimate we could get the worst and the best performance of the algorithm when compared to the other considered methods. The small system has the parameters *L* = 2, 10 initial molecules, and *k*_*d*_ = 10.0. The large system has the parameters *L* = 16, 10240 initial molecules, and *k*_*d*_ = 1.0. In the case of the large system, the methods marked with a red star were not able to finish the simulations because of system memory shortage (specific to the used machine) or for reaching the maximum computation time of 10 minutes per replicate. All results shown are an average of 10 replicates. In **(c)**,**(d)**,**(e)**, and **(f)** the computing time for varying number of reactions, number of initial molecules, diffusion rate *k*_*d*_, and total time of simulation are depicted respectively. The fixed parameters are *L* = 4, 100 initial molecules, *k*_*d*_ = 0.1, and total simulation time of 300 seconds. In **(g)** and **(h)** we plot the Principal Component Analysis for results obtained from simulations using 100 and 1,000,000 initial molecules in the system. Given that *L* = 4 used PCA to reduce the dimension of the vector of molecule counts *x* from 16 to 2 in order to be able to visualize the data. The variance explained in each axis is indicated in their respective labels.

### 8.2 Overall Performance Comparison

Given that the CBSA is designed to simulate large chemical systems, we first set two distinct scenarios using the diffusion model (a small and a large system) to compare the computational cost of the CBSA with other methods and implementations. The hypothesis is that the larger the system, the more impact the effect of matrix sparseness will have in reducing the CBSA computation time. For the small system case, we considered a system with *L* = 2, 10 initial molecules, and *k*_*d*_ = 10.0. For the large system we set *L* = 16, 10240 initial molecules, and *k*_*d*_ = 1.0. In Figure 5b we can see the mean computation time of 10 replicates for each of the considered methods in both small and large system cases. For the small system case, the sequential implementation of CBSA performed similarly to all SSA implementations except for STEPS. It is also observed that the *τ*-leaping implementations have no improvement of performance for such cases. On the other hand, the GPU implementation of CBSA obtained the worst performance for the small system. The reason for such limited performance is that GPU computations have a computational overhead due to memory transfers between host and device and also a lower clock speed. However, when the system is large enough, the efficiency gained with the computation distributed through the hundreds of GPU’s cores overcomes the memory transfer times making it marginal. This situation can be observed when considering the large system. Besides the higher efficiency of the sequential CBSA compared to the SSA methods, the gain in computation time is even better when using the GPU. For the large system, some methods were not able to conclude due to excessive use of the system’s memory or reached the maximum time established.

### 8.3 Parameter Scaling Comparison

To better understand how each variable of the diffusion model affects computation times, we performed simulations varying the square-lattice length *L*, the initial number of molecules, the diffusion rate *k*_*d*_, and the total simulation time. Regarding the system size, all the SSA implementations and the sequential CBSA showed an exponential growth in computing time as the value of *L* increased (Fig. 5c). On the other side, the GPU implementation of CBSA obtained a much lower growth rate in computational time as the size of the system grows, also performing better than all other methods when *L* ≥ 32. Similar exponential growth in computation time for all SSA implementations, except the StochPy Tauleap, was obtained when increasing the number of initial molecules in the system (Fig. 5d). However, the computational time is invariant to the number of molecular copies for both CBSA implementations as well as for the ODE solver.

It is expected in the SSA that the higher the reaction rates are, the more steps it takes to achieve a given simulation time. In Figure 5e we can observe that all SSA implementations present an exponential growth in the computation time as the diffusion constant is increased, with the *τ*-leaping implementations presenting a lower growth rate. The CBSA implementations only showed an increased computational time for the higher diffusion rate considered. This can happen when several reactions compete for the same reactant and the sum of their fluxes is higher than the available amount of the molecule. Thus, when several fast reactions share the same reactant, the CBSA needs to decrease the time step Δ*t* implicating in a higher number of steps to be computed in order to achieve a given simulation time. The variations in the time steps used in the CBSA is discussed in more detail in the Supplementary Information. For all methods, the computing time has a linear relationship with the total time of the simulation (Fig. 5f).

### 8.4 Some Observations about the Simulation

It is worth mentioning that one of the reasons for the increased performance of STEPS’ implementation in all the above-mentioned analyses is that it uses a limited number of random numbers generated before the actual simulation. One effect of this implementation choice can be observed when analyzing the variance of the molecular counts at the end of the simulations. In Figures 5g and 5h we can see a Principal Component Analysis (PCA) where the final molecular counts of all simulations are projected in a lower dimension space for two different number of initial molecules in the system. Considering that the simulation using ODE can be equivalent in some cases to the mean of several runs of a stochastic method, we can observe that in both PCA plots that the solutions provided by CBSA and SSA are scattered around the single solution provided by the ODE solver, with a lower scattering when the number of molecules in the system is higher. However, in Figure 5g we can observe only 2 out of 10 points simulated by STEPS because they are overlapping.

This result suggests that the limited number of random values has an impact on the stochasticity of the method, mostly when a lower number of molecules in the system is considered. It is also noticed that the solutions provided by the StochPy Tauleap are more scattered in Figure 5h than all other methods, which combined with the no variation in computation time might indicate an inconsistency in the solutions for a high number of molecules in this particular reaction system.

Although it is not the intent of this work to deliver a fully-featured software, both sequential and parallel implementations of CBSA, along with several other examples, are freely available at our GitHub repository^6^. The implementation and examples are also discussed in the Supplementary Information.

## 9 Concluding Remarks

Chemical reaction models are becoming more complex, especially when representing biochemical systems. Such models might currently involve up to thousands of molecular species and reaction channels becoming computationally challenging for current simulation methods. Also, some important characteristics should be appropriately handled on simulations of biochemical systems, such as the discreteness of molecular counts and the stochasticity of reaction occurrences.

Here we report a simulation method called Constraint-Based Simulation Algorithm – CBSA, which employs constraint-based modeling of chemical reactions in order to simulate large systems efficiently. We showed with theoretical examples that it could provide solutions similar to those produced by the most used method, the Stochastic Simulation Algorithm. It also yielded particularly good performance when simulating several system configurations, especially for a high number of molecules and reactions, thus fulfilling one of the initial motivations of obtaining a computationally scalable method.

Besides the benefits of CBSA already discussed in this work, we may point out other possible outcomes from this method. The so employed constraint-based modeling is widely used in the representation of biochemical systems. It should then be straightforward to apply CBSA on the simulation of virtually any model available in databases such as BiGG Models [33], and BioModels [34]. The practical implications of its use on real models should be assessed in future works. Furthermore, considering that the presented method was conceived with large biochemical models in mind, it can be a useful tool given the current advances in whole-cell modeling and simulation.

## Acknowledgements

PEPB thanks Bruno Messias and Thaãs G. Pedrosa for the insightful discussions regarding this work.

## Footnotes

1 All simulations were performed using the same computational architecture: Linux Debian 10 on an Intel i7-5930K, 64GB of DDR4 RAM, and NVidia GTX 1650 4GB.

2 StochPy: http://stochpy.sourceforge.net/

3 StochPy: http://http://steps.sourceforge.net/

4 GillesPy2: https://pypi.org/project/gillespy2/

5 pyOpenCL: https://pypi.org/project/pyopencl/

6 CBSA: https://github.com/pauloburke/CBSA

