## Supplementary Information for "Accelerated Simulation of Large Reaction Systems Using a Constraint-Based Algorithm"

### 1 Implementation of the Constraint-Based Simulation Algorithm

#### 1.1 Sparse-Matrix Implementation

In large chemical systems, it is likely that the representation using a constraint-based model will result in very sparse $S$ and $R*$ matrices, having a large amount of zero-valued entries. This means that the dot products involving these matrices will have to compute several pointless operations, leading to an unnecessary computational effort. To avoid such a situation and also reduce the memory usage of the system, we use a different representation of the $S$ and $R*$ matrices which will be described in more detail below.

Each step of the CBSA has two cost-wise major operations: the calculation of fluxes $v$ (Main Article Eq. 9) and the dot product $S \cdot v$ (Main Article Eq. 12 and 13). Both operations involve multiplications and/or sums through rows or columns. Thus, we want to reduce the number of operations by reducing the number of rows or columns of the matrices, removing as many zero-valued entries as possible. To do so, we will construct a substitution system for each of the two operations.

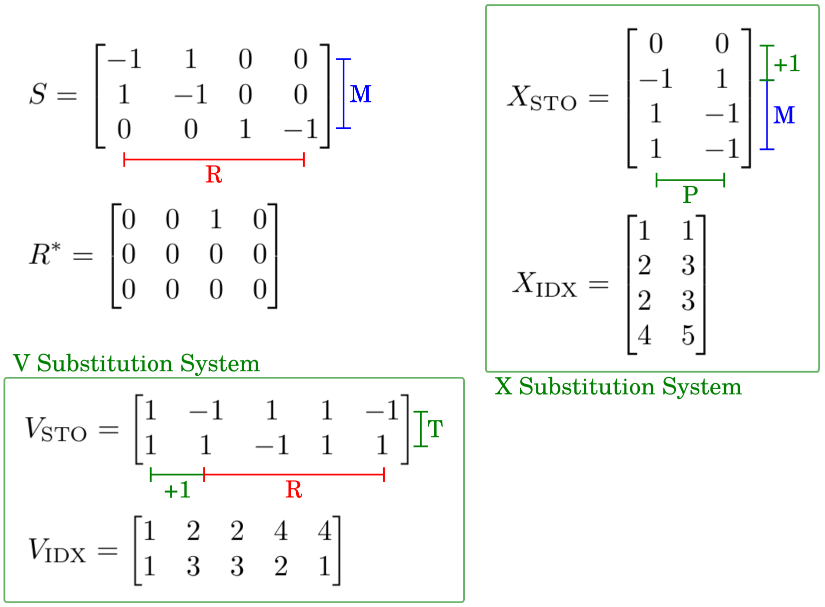

Figure S1: Example of the substitution system applied to the reaction system {1} of the Main Article. It is important to notice that the indexes in $V_{\mathrm{IDX}}$ and $X_{\mathrm{IDX}}$ are relative to the expanded vectors of $v$ and $x$. Thus, the indexes are increased by one relatively to the matrix $S$.

##### 1.1.1 Expansion of $x$ and $v$

The following substitution systems will only work after we expand the vectors $x$ and $v$ by inserting the values 0 and 1 at their respective initial positions, making their sizes $M+1$ and $R+1$ respectively.

##### 1.1.2 V Substitution System

To calculate $v$ in a more efficient way, let $V_{\text{STO}}$ and $V_{\text{IDX}}$ be two matrices of size $T \times R+1$ where $R$ is the number of reactions in the system and $T$ is the maximum number of reactants plus modifiers which a reaction in the system can have.

Each column in matrix $\boldsymbol{V}_{\text{STO}}$, starting at the second column, will be filled with the stoichiometry of the reactants and modifiers of the corresponding reaction. The entries not used in each column, as well as those from the first column, will be filled with the value one.

Each column in matrix $\boldsymbol{V}_{\text{IDX}}$, starting at the second column, will be filled with the indexes of the reactants and modifiers of the correspondent reaction. The entries not used in each column, as well as those from the first column, will be filled with the value one (or zero depending on the initial index assumed).

Then, we can rewrite the Equation 9 of the Main Article as:

$$v_i(t+\Delta t) = c_i \prod_{j=1,2,\ldots T} \frac{\binom{x_{V_{\text{IDX}ji}(t)}}{V_{\text{STO}ji}}}{V_{\text{STO}ji}!} \Delta t \tag{S1}$$

.

The new equation changes the number of multiplication operations from $2MR$ to $(R{+}1)T$. Given that $2M > T$, the number of operations will necessarily be reduced. Also, the greater the difference between $2M$ and $T$, the greater the gain obtained by applying the V substitution system.

##### 1.1.3 X Substitution System

Similarly to the method presented in the last section, we will build a substitution system to calculate $S \cdot v$. Let $X_{\text{STO}}$ and $X_{\text{IDX}}$ be two matrices of size $M \times P+1$ where $M$ is the number of molecules in the system and $P$ is the maximum number of reactions a molecule participates in the reaction system.

Each line in matrix $\boldsymbol{X}_{\text{STO}}$, starting at the second one, will be filled with the stoichiometry in reach reaction that the correspondent molecule participates. The entries not used in each row, as well as those from the first row, will be filled with the value zero.

Each line in matrix $\boldsymbol{X}_{\text{IDX}}$, starting at the second row, will be filled with the indexes of the reactions that the correspondent molecule participates. The entries not used in each row, as well as those from the first row, will be filled with the value one (or zero depending on the initial index assumed).

Then, we can rewrite $\Delta x = S \cdot v$ as follows:

$$\Delta x_i = \sum_{j=1,2,\ldots P} X_{\text{STO}ij} v_{X_{\text{IDX}ij}} \tag{S2}$$

.

The new equation changes the number of multiplication and sum operations from $MR$ to $(M+1)P$. Given that $R \geq P$, the greater the difference between $R$ and $P$, the greater the gain obtained by applying the X substitution system.

### 2 Examples

#### 2.1 Two Reactions Model

Let us consider the set of two reactions described in {S1}. The molecule A can be converted in a molecule B or C with reaction rates $c_1$ and $c_2$ respectively.

$$\begin{aligned} A &\xrightarrow{c_1} B \\ A &\xrightarrow{c_2} C \end{aligned} \tag{\{S1\}}$$

Figure S2 depicts several trajectories obtained by simulating this system using the CBSA. A very similar result is obtained using the SSA. The histogram under the plot shows the distribution of the time at which the number of molecules A reaches zero as it is consumed by both reactions. The histograms on the left show the distribution of the number of B and C molecules after 15 seconds.

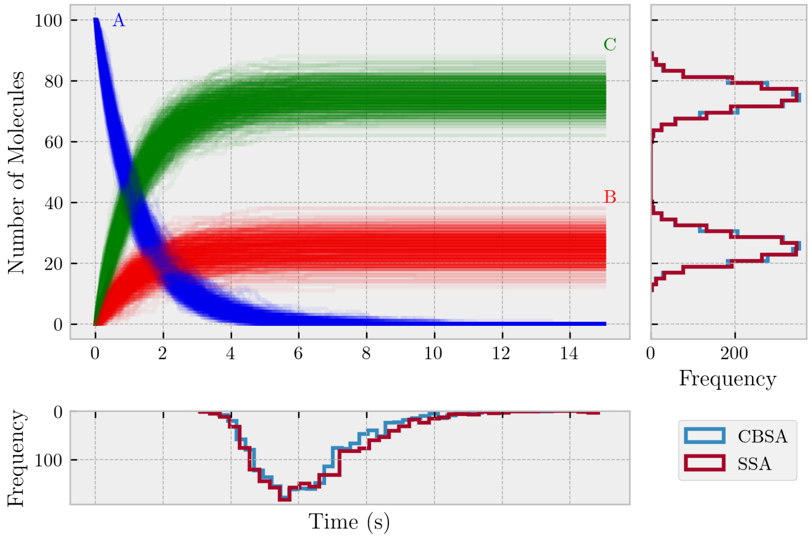

Figure S2: Simulation of the chemical system described in {S1} using the CBSA.

#### 2.2 N-Reactions Model

The CBSA first calculates the reaction fluxes as if they were independent. However, if two or more reactions share the same reactants, they can compete for them and their fluxes will no longer be dependent. Thus, if the amount requested of a given molecule by the reactions is bigger than the available amount, the set of calculated fluxes will not pass the mass-balance constraint. In such a case, the algorithm will search for a $\Delta t$ that when multiplied by the fluxes will satisfy the constraint.

The search for valid fluxes is accomplished by multiplying the current $\Delta t$ by a constant $0 > \alpha < 1$ until $v\Delta t$ meets the mass-balance constraint. The number of times that the multiplication by $\alpha$ is necessary in order to find a valid set of fluxes depends on the system and the system‘s current state, impacting the time demanded by each step in the simulation.

To better understand the impact of $\Delta t$ and $\alpha$ on the simulations using the CBSA, consider the chemical system described in S2. The system contains $n+1$ reactions where the first is a spontaneous creation of molecule A at a rate $c_0$ and the other $n$ reactions will convert A into $B_1, B_2, ...B_n$ molecules with a rate $c_1$. For some combinations of $n$, $c_0$, $c_1$, and $\Delta t$ the reactions will compete for A, leading the algorithm to search for a a valid $v\Delta t$.

$$\begin{aligned}
\varnothing &\xrightarrow{c_0} \mathrm{A} \\
\mathrm{A} &\xrightarrow{c_1} \mathrm{B_1} \\
\mathrm{A} &\xrightarrow{c_1} \mathrm{B_2} \\
&... \\
\mathrm{A} &\xrightarrow{c_1} \mathrm{B_n}
\end{aligned} \qquad \{S2\}$$

We simulated the system described in {S2} for different values of $n$. We also considered different values of $\Delta t$ and $\alpha$. Figure S3 shows some statistics from 50 simulation replicates for each combination of $n$, $\Delta t$, and $\alpha$. Overall, we can observe that as $n$ grows, the competition for A is intensified, implying a decreasing mean step length from the initial $\delta t$. It also implies a higher computational cost.

The reduction in the step length is due to the multiplications of $\Delta t$ by $\alpha$ to find a valid solution. In Figure S4 we can see how many $\alpha$-iterations the algorithm takes to find a valid solution given different initial $\Delta t$ and $\alpha$. As $n$ grows, the number of $\alpha$-iterations in each step of the algorithm increases, abbreviating the time-step taken. Therefore, more steps are required to achieve the desired stopping criterion.

Regarding the error in the simulation outputs, we compared the final number of $B_1, B_2, ...B_n$ molecules with an expected theoretical mean value. Except when $\Delta t = 1.0$, for almost all other combinations of parameters the error remained below 1%. To analyze the standard deviation of the final values, we compared them to the ones obtained with the SSA. Excepting when $\Delta t = 1.0$, all standard deviations obtained presented an error lower than 10%.

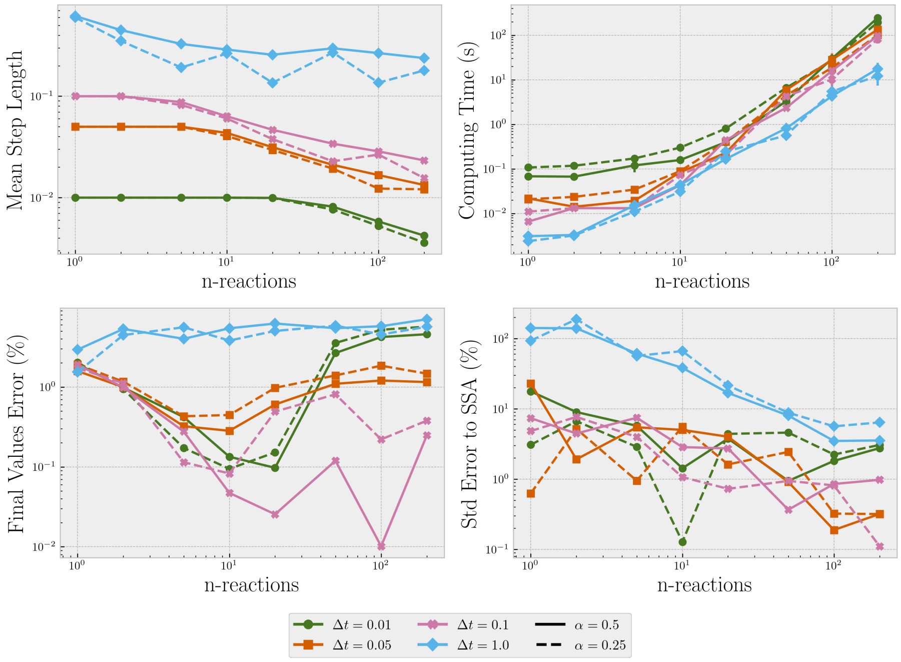

Figure S3: Analysis of mean step length, computing time, and relative errors from 50 simulation replicates of the system {S2} for different number $n$ of reactions. The stopping criterion was set as 10 seconds. The errors for both final values and standard deviations were calculated using the root-mean-square error (RMSE).

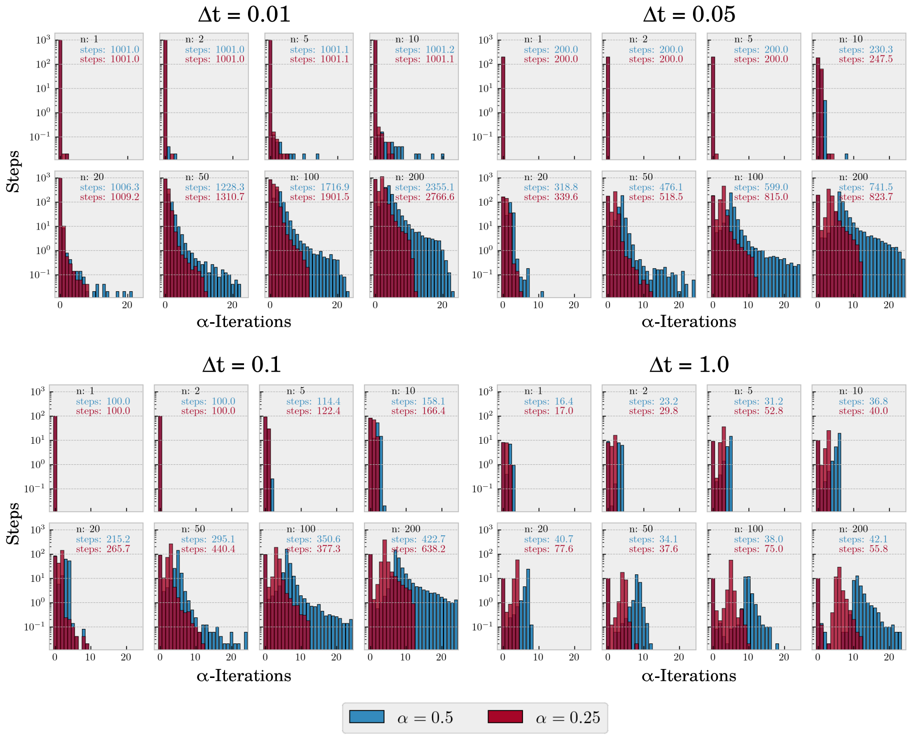

Figure S4: Histograms of the number of $\alpha$-iterations that each step in the algorithm takes to find a valid solution.

#### 2.3 The Oregonator

In the original publication of the Stochastic Simulation Algorithm (SSA) by Daniel T. Gillespi [1], one of the models used to show the algorithm's capabilities was the Oregonator. This model is a chemical oscillator with closed limit cycles proposed by Field and Noyes [2] and is composed as follows:

$$\begin{aligned} X_1 + Y_2 &\xrightarrow{\mathrm{c}_1} Y_1 \\ Y_1 + Y_2 &\xrightarrow{\mathrm{c}_2} Z_1 \\ X_2 + Y_1 &\xrightarrow{\mathrm{c}_3} 2Y_1 + Y_3 \\ 2Y_2 &\xrightarrow{\mathrm{c}_4} Z_2 \\ X_3 + Y_3 &\xrightarrow{\mathrm{c}_5} Y_2 \end{aligned} \tag{\{S3\}}$$

.

Given that the number of $X_1$, $X_2$, and $X_3$ will be constant along time, we can rewrite the equations transferring their values to the reactions rates:

$$\begin{aligned} Y_2 &\xrightarrow{\mathrm{c}_2\mathrm{X}_1} Y_1 \\ Y_1 + Y_2 &\xrightarrow{\mathrm{c}_2} \varnothing \\ Y_1 &\xrightarrow{\mathrm{c}_3\mathrm{X}_2} 2Y_1 + Y_3 \\ 2Y_2 &\xrightarrow{\mathrm{c}_4} \varnothing \\ Y_3 &\xrightarrow{\mathrm{c}_5\mathrm{X}_3} Y_2 \end{aligned} \tag{\{S4\}}$$

.

In the same way as performed in [1], we rearranged the reaction rates as a function of the initial amounts of $Y_1$, $Y_2$, and $Y_3$ as well as two additional parameters $\rho_1$ and $\rho_2$:

$$\begin{aligned} c_1 X_1 &= \rho_1 / Y_{2s} \\ c_2 &= \rho_2 / Y_{1s} Y_{2s} \\ c_3 X_2 &= (\rho_1 + \rho_2)/Y_{1s} \\ c_4 &= 2\rho_1 / Y_{1s}^2 \\ c_5 X_3 &= (\rho_1 + \rho_2)/Y_{3s} \end{aligned} \tag{S3}$$

.

The reaction system described in {S4} can be written as a constraint-based model in terms of $S$ and $R*$ as follows:

$$S = \begin{array}{c} \begin{matrix} r_1 & r_2 & r_3 & r_4 & r_5 \end{matrix} \\ \begin{bmatrix} 1 & -1 & 1 & -2 & 0 \\ -1 & -1 & 0 & 0 & 1 \\ 0 & 0 & 1 & 0 & -1 \end{bmatrix} \end{array} \begin{matrix} Y_1 \\ Y_2 \\ Y_3 \end{matrix} \tag{S4}$$

$$R^* = \begin{array}{cc} \begin{matrix} r_1 & r_2 & r_3 & r_4 & r_5 \end{matrix} & \\ \begin{bmatrix} 0 & 0 & 1 & 0 & 0 \\ 0 & 0 & 0 & 0 & 0 \\ 0 & 0 & 0 & 0 & 0 \end{bmatrix} & \begin{matrix} Y_1 \\ Y_2 \\ Y_3 \end{matrix} \end{array} \tag{S5}$$

.

In Figure S5 we can observe that the CBSA can reproduce the results shown in [1]. The system presents a well-defined limit cycle after a small period of convergence.

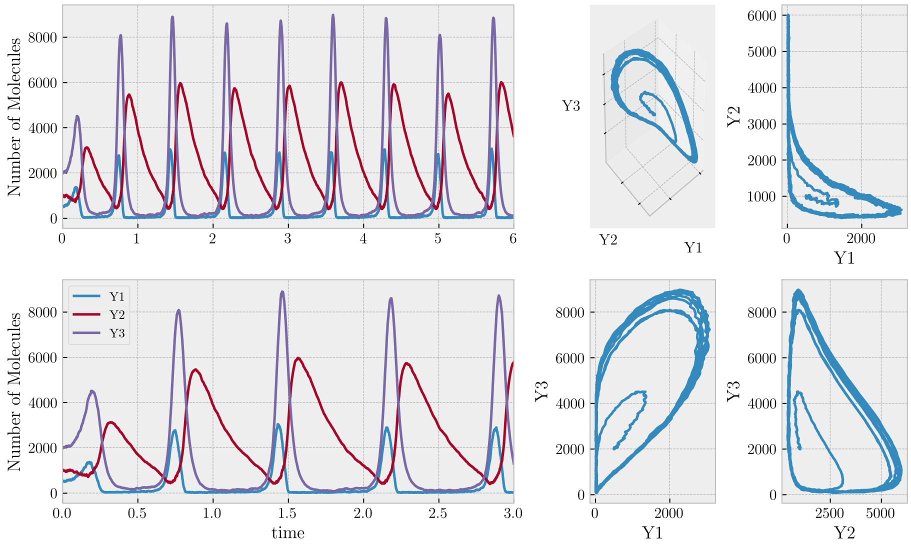

Figure S5: Simulation of the Oregonator using the CBSA with the following parameters: $Y_{1s} = 500$,$Y_{2s} = 1000$,$Y_{3s} = 2000$, $\rho_1 = 2000$, $\rho_2 = 50000$, $alpha = 0.5$, and $\Delta t = 0.001$.

#### 2.4 Bi-Stable Genetic Switch

One of the goals of Synthetic Biology is to modify living organisms in order to better control their behavior. In 2014, Jerala *et al.* managed to insert an artificial genetic circuit into a human cell so that it could produce two different fluorescent proteins as a response to the presence of two particular activator molecules in the environment [3]. Additionally to the response to the signal, the cells could also retain the memory of their state even when the activator molecule was no longer in the medium or change their state if the signal molecule was changed. The state of the cells could be visually checked through fluorescence microscopy.

The kind of behavior described above can be called a bi-stable switch. The system has two stable states and can transit between each other given a respective stimulus. It was implemented in [3] by employing positive and negative feedback loops in the expression of genetic constructs that contain the genetic code for receptors to signal molecules, promoter enhancers and inhibitors, and the fluorescent reporter molecules. They also provided a computational model of their genetic circuit as well as parameters obtained experimentally.

Here we reproduce the simulations of the bi-stable genetic switch model using the CBSA. The set of reactions and reaction rates were obtained from Tables 15 and 7 respectively from the Supplementary Material of [3]. The model comprises a total of 28 molecular species and 46 reactions.

In Figure S6 we can see how the levels of the GFP and mCitrine (mCT) reporter molecules change in different scenarios, with or without the presence of the pristinamycin (PI) or erythromycin (ER) inducer molecules. Each curve corresponds to an independent simulation which can be interpreted as an individual cell.

The results obtained using the CBSA are consistent with the simulations and experimental data available in [3]. The presence of PI and ER induces the production of BFP and mCT respectively. The removal of the inducers after a certain period does not affect the production of the respective reporters evidencing the system's memory. The change of ER to PI triggers a change of state in the cell which gradually stops to produce mCT and begins to produce BFP.

It was shown experimentally in [3], but not considering simulations, that cells treated with no inducer tend randomly to one of the two states due to random fluctuations in the expression rates of enhancers and inhibitors. The same behavior could be observed in the simulations shown in Figure S6.

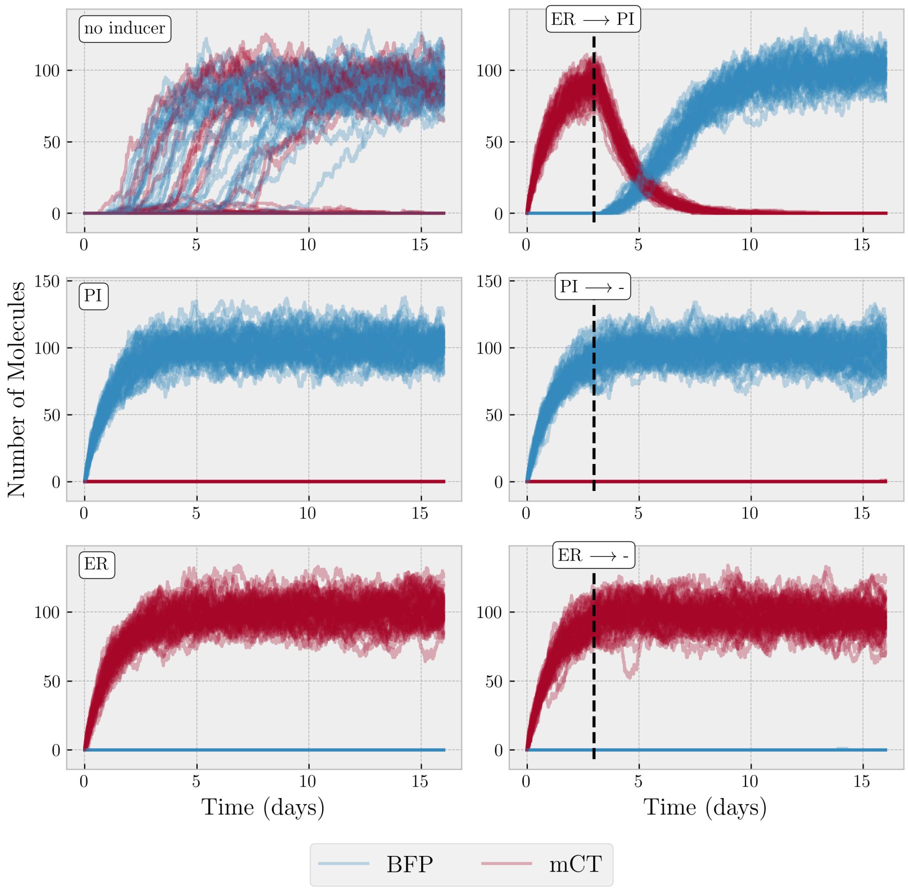

Figure S6: Simulations of the bi-stable genetic switch using the CBSA. The black dashed lines indicate the moment when the inducer present in the environment was changed or removed. Each curve corresponds to one of 50 independent simulations, each one taking approximately $17 \pm 3$ minutes to compute using $\alpha = 0.5$ and $\Delta t = 1.0$ seconds.
